# piRNAs from Y chromosomal protein coding, noncoding and endogenous retrovirus homologous repeat families regulate autosomal gene expression in mouse testis

**DOI:** 10.64898/2026.06.23.733120

**Authors:** Rachel A Jesudasan, Arpan Mukhoti, Anurag Chaturvedi, Shrish Tiwari, Kankadeb Mishra, Pranatharthi Annapurna, Nissankararao M Praveena, Jomini L Alex, Sivarajan Karunanithi, Arvind Kumar, Hemakumar M Reddy

## Abstract

**Background:** The heterochromatic long arm of the mouse Y chromosome harbors multicopy species-specific sequences *Ssty, Sly, Asty* and *Orly* that are transcribed in testis and have known functions in male fertility. Multiple isoforms have been described for each of these. Previous studies reported some isoforms of *Ssty* and *Sly* as coding and others as noncoding. Here we investigate further, the roles of these Y-heterochromatin transcripts. Earlier proteomics studies identified a few proteins which were expressed differentially between wild type (XY^RIII^) and Y-deletion mutant (XY^RIII^qdel) mice. These genes were regulated by piRNAs derived from a noncoding Y-transcript, *Pirmy*. The multicopy Y-transcripts were analyzed for similar regulatory functions in this study.

**Methods:** XY^RIII^ and XY^RIII^qdel mice were used. Bioinformatics analysis and small RNA northern blots predicted and identified piRNAs from these Y-transcripts. Luciferase reporter assay indicated functional relevance. dPCR, RT-qPCR and western blotting using XY^RIII^ and XY^RIII^qdel mice confirmed regulation of autosomal genes.

**Results:** Multicopy gene families from mouse Y-heterochromatin generate piRNAs that putatively regulate six autosomal genes. Additionally, the ERV-LTR, found in the *Orly1* transcript had homology to UTRs of a few autosomal genes and identified piRNAs in the piRBase database. *Orly1* showed a reduction in genomic copy number in XY^RIII^qdel mice. *Pm20d1*, one of the four autosomal genes harboring the ERV-LTR segment in its UTR, showed upregulated protein expression in XY^RIII^qdel testes.

**Conclusions:** Thus, we demonstrate that different classes of repeats from Y-chromosome regulate autosomal gene expression via piRNAs. In addition, this study also identified a novel role for a Y-derived ERV in autosomal gene regulation in testis.

## 1. Introduction

The role of mammalian Y chromosomes in testis-determination, spermatogenesis and male fertility is well established [1-4]. Y-chromosomes in general contain a few protein-coding genes and are replete with repeats [5-7]. The entire long arm of the mouse Y-chromosome stains heterochromatically and harbors a unique repertoire of coding and non-coding, sex and species-specific multi-copy genes that are expressed in testis [8-10]. The testis-specific *Ssty* (spermiogenesis-specific transcript on the Y), *Sly* (*Sycp3*-like Y-linked), *Asty* (amplified spermatogenic transcripts Y encoded) and *Orly* (oppositely transcribed, rearranged locus on the Y), are multicopy genes present on the long arm of the mouse Y chromosome [11-14].

Whereas *Ssty* and *Sly* families code for proteins, *Asty* and *Orly* transcripts do not appear to have coding potential [8]. *Orly* comprises regions from *Sly, Ssty* and *Asty*, three known Yq-linked gene families. Large numbers of *Orly* isoforms are perhaps transcribed at low levels [8]. The terminal exon of *Orly1* is derived from a partially degenerate Murine Repeated Virus on the Y chromosome (MuRVY) [8], a class 1 endogenous retroviral element (ERV). *Orly1* does not have a significant ORF other than a partial degenerate retroviral Pol sequence and thus is considered noncoding. *Orly* is transcribed testis-specifically [8]. Approximately 10% of the mouse genome harbors Endogenous Retroviral Elements (ERV) [15]. There are multiple copies of *Ssty* with two distinct intronless subfamilies, *Ssty1* and *Ssty2* which code for SSTY1 and SLY proteins [10, 13]. that are expressed in mouse spermatids [11]. SLY is thought to be involved in multiple processes in spermiogenesis as it interacts with an acrosomal protein, a histone acetyl transferase and a microtubule-associated protein. Although some copies of *Ssty1, Ssty2* and *Sly* code for proteins, others are transcribed but not translated [4, 10, 11, 14].

Mice with Y chromosome long arm deletions show morphological and motility related abnormalities in sperm, and exhibit either subfertility/sterility depending on the extent of the deletion [10]. Mice with 2/3^rd^ deletion of the long arm of the Y-chromosome (XY^RIII^qdel) exhibit sperm head abnormalities, sperm motility aberrations, reduced sperm count, skewed sex ratio towards females in the progeny and subfertility [4]. These deletions also lead to upregulation of several X- and Y-transcripts in spermatids [12]. The proteins encoded by the *Ssty* and *Sly* gene families were earlier considered to be responsible for the phenotypes observed in mice with deletions of Yq [10, 11].

Functional significance of the noncoding transcripts is not known. Earlier studies revealed UTR-mediated regulation of several genes in testis and brain by Y-heterochromatin-derived noncoding transcripts in mouse and man [9,16,17]. The *Pirmy* and *Pirmy*-like transcripts described from mouse Y-heterochromatin are noncoding transcripts which regulate several autosomal genes via piRNAs. A comparative proteomics screen between XY^RIII^ and XY^RIII^qdel identified proteins that are differentially expressed. These autosomal genes are regulated by *Pirmy*-derived piRNAs. [9]

In the current study piRNAs derived from the Y-transcripts analyzed here, are shown to regulate some of the differentially expressed proteins in the mutant mice. ERV-LTR element from *Orly1* present in the UTRs of five autosomal genes putatively regulate these genes via piRNAs. Thus, in the present study, we provide experimental evidence (1) to support generation of piRNAs from protein coding and noncoding repeat gene families localized on the mouse Y long arm and (2) that they regulate the autosomal genes.

## 2. Materials and Methods

XY^RII^ and XY^RIII^qdel mice [18] used were obtained as a gift from Prof. Paul S Burgoyne and bred in our animal house facility in accordance with the guidelines from Indian Science Academy under CPCSEA (Committee for the Purpose of Control and Supervision of Experimental Animals). The tissues (brain, kidney, testis, ovary, heart and spleen) used in the small RNA Northern blots, i.e., were collected from 2-3 months old XY^RIII^ and XY^RIII^qdel mice. Freshly isolated tissues were flash frozen in liquid nitrogen and stored in -80 ° C freezers till required. We isolated small RNA from these tissues, pooled to obtain sufficient quantity, and transferred them on to Hybond N+ membrane for northern blot analysis as per protocols described earlier [9]. At least three biological replicates were done to confirm the results obtained.

### Electrophoretic Mobility shift Assay (EMSA)

EMSA was done with MIWI protein following published protocols [9]. RNA oligonucleotides were synthesized from the UTR homologous regions of *Ssty* (5’CCUCAUGAAGAAGAGGAGGAGGA3’ 476-498) and *Sly* (5’CAGUUAAAGAAAUGCAUGAGAA3’ 666-687) for EMSA. piR-mmu-1, a known piRNA (5’UGACAUGAACACAGGUGCUCAGAUAGCUUU3’) [19], was used as a control. Radio-labelled oligos and purified MIWI protein in a reaction buffer standardized earlier in the lab, were used for checking the binding of putative piRNAs to MIWI. Unlabeled oligos and a known piRNA, piR-mmu-1 were used as cold competitors. MIWI (G82) antibody (90ng, Cell Signaling Technology, Cat # 2079S) or Argonaute 3 antibody (100 ng, Abcam, Cat # ab3593), were procured from the vendors specified here. EMSA was done on 5% native PAGE and image captured using FUJI phosphorimager (FUJIFILM FLA-3000). Argonaute 3 antibody served as the antibody control. At least three biological replicates were done to obtain consistent results.

### Luciferase Assay

This assay was done using the 5’ UTR of RP11-130C19 clone. We assayed the reporter gene expression with anti-oligomers named as antagopirs and the 5’UTR. Figure 2a is a schematic representation of the 5’UTR reporter construct. In silico analysis using *Ssty2* sequence identified a homologous stretch in the 5’UTR of the gene that localized to the clone RP11-130C19. Complimentary RNA oligo nucleotide was procured from Eurofins Genomics Pvt. Ltd., Bangalore, India. Briefly, the 5’UTR was cloned into pSP_luc+NF fusion vector transcript (Promega Cat #E4471). Luciferase gene alone and the Luciferase gene with UTR were cloned separately in pcDNA 3.1 expression vector (Invitrogen, Cat # V790_20), for assaying the effect of antagopirs on Luciferase expression. GC_1spg cell line (ATCC LRL-2053), derived from mouse testis, and lipofectamine 2000 Transfection Reagent (Thermo Fisher Scientific Cat # 11668019) were used in the transfection experiments using previously published protocols [9]. The transfection and reporter assays were done as described earlier [9]. Co-transfection with 50ng of β− gal plasmid was done to assess transfection efficiency. At least three biological replicates were performed. This included three technical replicates in each experiment. Statistical analysis was done to check for variability and significance.

**Figure 1.**
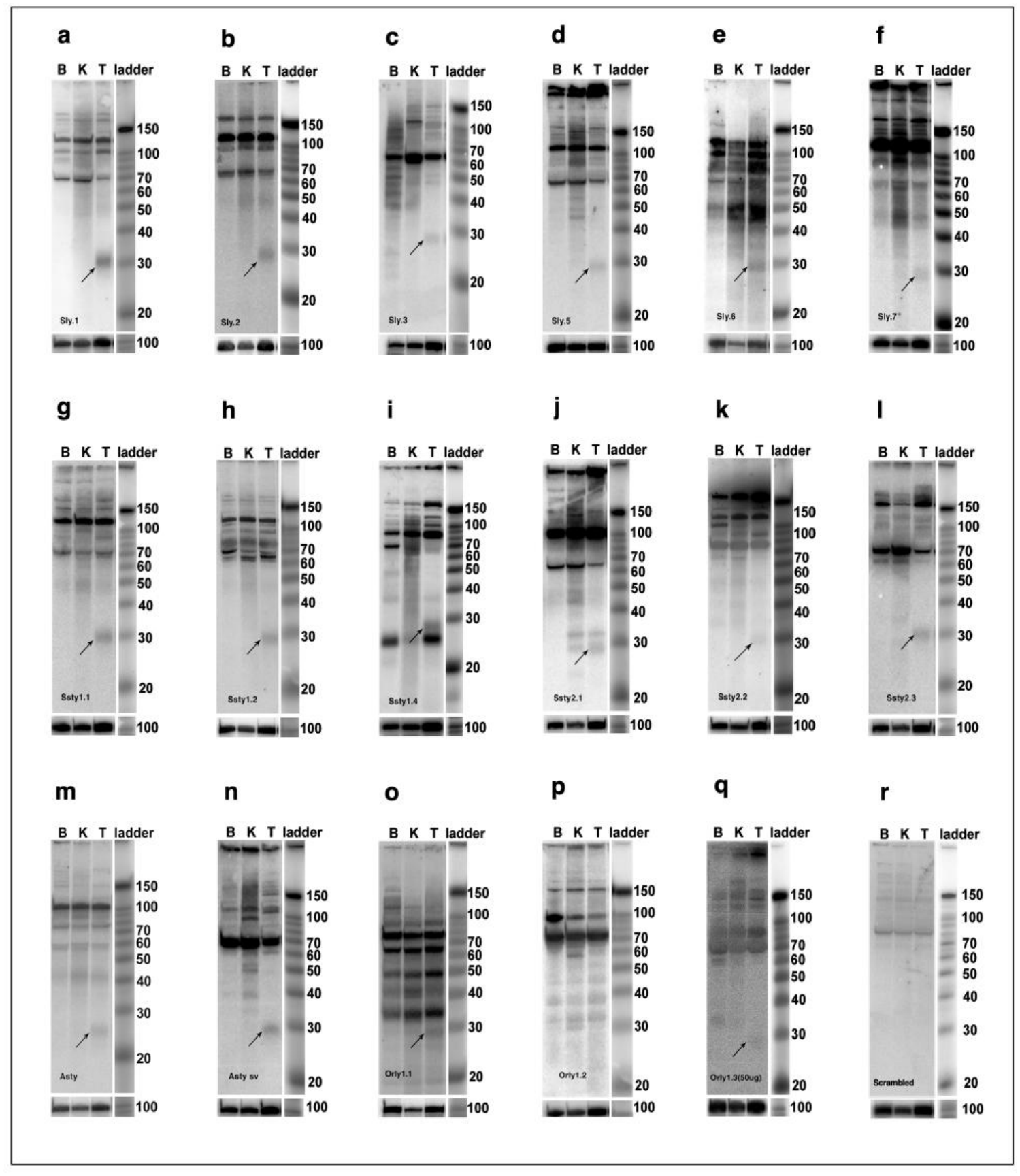
Small RNA northern blots showing signals corresponding to the size of piRNAs: LNA oligonucleotides showing homology to Y-chromosomal transcripts and UTRs of different genes - were used as probes on northern blots containing 20 µg small RNA from brain (B), kidney (K) and testis (T). Testis-specific signals of 29-30 nt observed with all *Sly* probes (a-f).LNA probes from *Ssty1***(g-i)**; *Ssty2* **(j-l)** and *Asty* (**m**), *Asty-sv* **(n)** show signals ranging from 26-32 nt. Signals from *Orly1* are shown in sub panels (**o-q)**. A scrambled probe was used as the negative control **(r)**.

**Figure 2.**
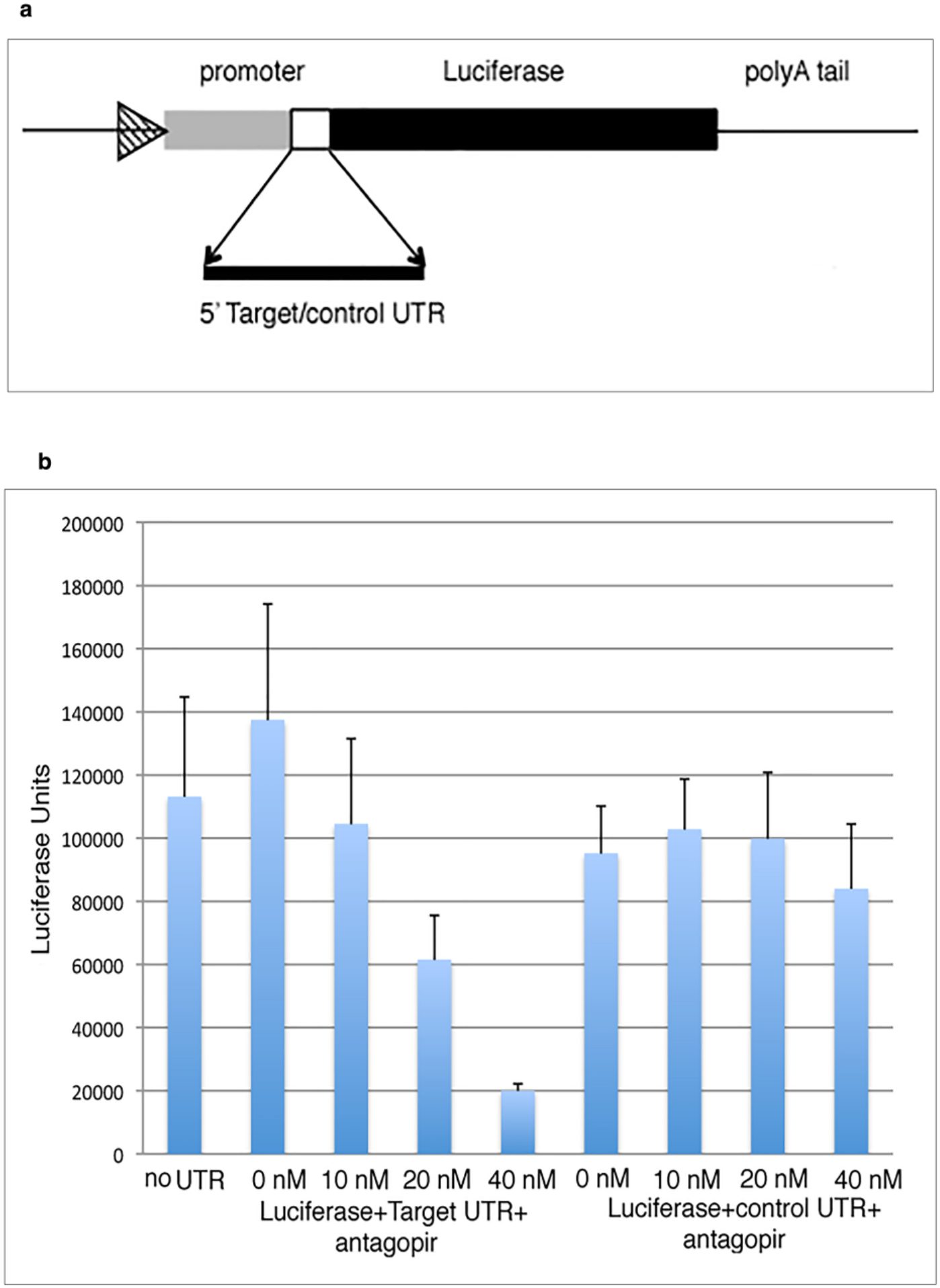
Reporter assay for functional analysis of piRNAs. (**a**) Cartoon representing Luciferase construct, where 5’UTR (RP11-130C19 gene/ *Gapdh*) was inserted between promoter and coding region of Luciferase gene; (**b**) Histogram representation of luciferase assay results from control and target UTRs containing homology to Y-transcripts with different concentrations of antagopir. Figure shows decrease in Luciferase levels with increasing concentrations of antagopir using the target UTR construct, whereas with the control UTR construct *(Gapdh)*, antagopir does not show a concentration dependent reduction in luciferase expression.

### piRNA PCR analysis

Tissues collected from male and female mice, of about 2-3 months of age of the C57/BL6-Ncrl strain and from XY^RIII^ and XY^RIII^qdel mice were used. Animals were sacrificed by cervical dislocation and collected tissues were flash-frozen in liquid nitrogen. After isolation of small RNA, equal amount from 3 animals were pooled to a final concentration of 30ng/µl. The standard protocol provided with the miRCURY LNA RT-Kit from Qiagen (Catalogue No: 339340) was followed for cDNA synthesis, with slight modifications. A maximum reaction volume of 10µl was found to be optimal, as increasing the reaction volume lowered efficiency. When a larger amount of cDNA was required, multiple reactions were set up. Also, 60ng of RNA, instead of 10ng as suggested in the protocol, was found to be more optimal. qPCR reactions were set up in QuantStudio 3000 system, using standard SYBR Green-based method with TB Green II by Takara Bioscience (Catalogue No: RR820). piRNA-specific primers were used as forward primers, and a Locked Nucleic Acid (LNA) oligo (dT) complementary to the poly A tail attached to the small RNA, was used as a reverse primer. piR-mmu-1 was used as a positive control and miR-2 was used as an internal control, for the qPCR reactions. The reactions were carried out for 40 cycles. ΔΔct analysis was used to calculate the expression levels of piRNAs. Statistical significance was calculated using Students t-test.

### Statistical Analysis

The two tailed unpaired student’s t test (Welch’s t test) was employed to determine statistical significance between the XY^RIII^ and XY^RIII^qdel groups in the piRNA RT-qPCR studies and the western blots. The significant p values were expressed as follows: *= p≤ 0.05, **=p≤ 0.01, ***= p≤ 0.001.

### Computational Analysis

A BLAST analysis using default parameters, of the 8 Y transcripts (*Orly1, Orly2, Orly3, Sly, Ssty1, Ssty2, Asty, Asty-sv*) against the UTRs of mouse (from the UTRdb; the GRCm37 assembly) was performed. The hits, with e-value < 0.01, were further analyzed using experimental and Bioinformatics approaches. For identification of piRNAs from predicted motifs, BLAST analysis was done against different piRNA databases including the Gold standard piRNAs. Gold standard piRNA sequences for six species—*Homo sapiens* (Hsa), *Drosophila melanogaster* (*Dme*), *Mus musculus* (*Mmu*), *Macaca fascicularis* (*Mfa*), *Rattus norvegicus (Rno)*, and *Bos taurus (Bta)*—were downloaded from piRBase release v3.0 (http://bigdata.ibp.ac.cn/piRBase/download.php). The gold standard set represents a smaller, more representative subset of piRNAs selected according to piRNA characteristics from the tens of millions of sequences in the database. *Orly2* and *Orly3* did not identify piRNAs. A nucleotide blast (blastn; v2.10.1) was therefore performed using the 6 Y transcripts (*Sly, Ssty1, Ssty2, Asty, Asty-sv, Orly1*) against the gold standard set, with the word size set to 7 to increase sensitivity and limited the results to the top 10 best-scoring alignments per query (max_target_seqs 10). All other parameters were set to default.

### dPCR for copy number estimation

Mouse spleen and gonads were used for isolation of genomic DNA using standard protocols. For absolute quantification of genomic copy number of *Sly* and *Orly* genes, we used the QIAcuity EG PCR kit by QIAgen. DNA samples were diluted to a concentration of 20 ng/µl, and further diluted 100x before setting up the reactions. Each reaction was set up in a total volume of 40µl in a well of QIAquity Nanoplate 26k-24 wells. 2.5µl of diluted DNA (500 pg) was used, following the QIAgen protocol. PCR was done under the following cycling conditions - 15s of denaturation at 95°C; 15s of annealing at 54°C; 15s of Extension at 72°C for 40 cycles. Readings were taken by recording the fluorescence at an exposure of 100ms. To calculate the absolute number of copies of a gene per cell we first calculated the number of copies/µl (cp/µl) in the stock DNA from the cp/µl of the reaction mix by the following formula - (Cp/µl X 40)/2.5) X 100). We calculated the number of copies present in 1 ng of DNA by the following formula (Cp/µl in stock DNA)/ (concentration of stock DNA in ng/µl). This was then multiplied by 6×10^-3^ to calculate the number of copies per cell (6 pg is the approximate weight of total chromosomal DNA in a single cell).

## 3. Results

### 3.1. Transcripts from multicopy gene families on mouse Yq generate piRNAs in testis

BLAST analysis of the transcripts derived from multicopy genes on mouse Y chromosome viz., *Sly, Ssty1, Ssty2, Asty*, and *Orly* against the UTR database (based on assembly NCBI37/mm9) led to the identification of short (11-30 nucleotides) and long stretches of homology (>150 nt), in 3’ and 5’ UTRs of many autosomal genes in both +/+ (direct) and +/- (complementary) orientations using default parameters. Number and orientation of short hits for each Y-transcript is shown in (Table S1).

#### 3.1.1. Experimental validation of piRNA

##### 3.1.1.1. Small RNA northern blot analysis

Locked nucleic acid (LNA) oligonucleotide probes (11-23 nt) homologous to different UTRs that have matches to the Y-transcripts were used as probes on small RNA blots (20 µg in each lane). Many Y-chromosomal sequences have homologous X-counterparts. The probes were checked for homology to mouse X chromosome to eliminate such sequences (e-values <0.01). Only the probes specific to Y were chosen for northern blot analysis. Sixteen of the seventeen probes identified 26-32 nt long signals predominantly in testis, which suggested that these could be piRNAs based on their size (Figure 1a-q); Most of the probes identified signals corresponding to piRNAs with 20 µg small RNA on northern blots. Orly1.3 did not identify testis-specific signals even with 30 µg of small RNA. Orly1.3 identified a faint signal in testis, on a 50 µg-small RNA blot (Figure 1q) - indicating low levels of expression of these transcripts/piRNAs. We observed piRNAs mainly in testis although two of the oligonucleotides identified signals, corresponding to piRNAs in brain and kidney (Fig. 1i and j). A scrambled probe that was used as negative control did not give any signal in the range of small RNAs (∼26-32 nucleotides) in any of the tissues, (Figure 1r) indicating that the signals observed with the probes were specific. A probe corresponding to U6, which is a highly conserved, ubiquitously transcribed, low molecular weight small nuclear RNA served as the loading control for all the blots (Figure 1a-r). To demonstrate that these small probes did not identify nonspecific signals, two of the Y-derived 11 nucleotide long probes Ssty1.2 and Sly.1 were used to examine expression in other tissues such as the male heart, spleen and the female brain, ovary. The probes identified signals only in the testis (Figure S1a, b), confirming specificity of the probes. The localization of probes used for small RNA northern blotting is depicted on the corresponding Y-transcripts (Table S2; Figure S2).

Computational analysis of the primary transcripts, *Sly, Ssty1, Ssty2, Asty, Asty-sv* and *Orly1* did not reveal any stable secondary structure, on using sliding-window technique with window size of 70 nt and window position increment of 1 nt (data not shown), using pknotsRG [19]. These transcripts do not have stable secondary structures that are characteristic of miRNAs. BLAST analysis of the Y-transcripts against the mouse gold standard piRNAs from piRBase database (https:/bigdata.ibp.ac.cn/piRBase) identified piRNAs, indicating that these homologous stretches could be piRNAs. Table S3.

##### 3.1.1.2. Electrophoretic Mobility Shift Assay

The *Sly* and *Ssty* probes showed shift in mobility upon incubation with purified MIWI protein (Figure S3a and b). The mobility shift was competed out with either cold piR-mmu-1, a known piRNA [19] or MIWI antibody. Similarly, cold *Ssty* and *Sly* oligonucleotides as well as the MIWI antibody competed out binding of piR-mmu-1 to MIWI protein i This indicated that *Ssty, Sly* derived oligonucleotides were piRNAs. When an alternate antibody, argonaute 3 antibody, was used as a competitor in the gel shift assays, it did not compete out the binding of MIWI protein to the oligonucleotides. The results obtained from EMSA were consistent with the possibility that the studied sequences are piRNAs.

##### 3.1.1.3. Y-derived piRNAs are expressed differentially from the two strands of DNA

Oligonucleotide probes Sly.4, Ssty1.4 and Orly.1 designed forboth direct (sense, D) and complementary strands (antisense, C) (Figure S4) yielded signals corresponding to the size of piRNAs with sense probes only, under identical conditions, revealing differential expression of piRNAs from the two strands (Figure S4a-for *Sly* only, others data not shown). However, stringent washing and processing the blots in the sigmoid scale, elicited∼30 nt-long signals in testis with antisense probes also (Figure S4b-for *Sly* only, data not shown for other probes). Thus, both the sense and antisense strands of DNA produce piRNAs – albeit in unequal quantities.

#### 3.1.2. Functional analysis of piRNA sequences

Regulation of gene expression by some of these piRNAs were studied using the Luciferase assay. *SSty2* transcript has homology to the 5’UTR of the clone RP11-130C19. Increasing concentrations of the antagopir, i. e. 10 nM, 20 nM and 40 nM caused concentration dependent reduction in Luciferase expression (Figure 2). The antagopir did not have an effect when the UTR from the *Gapdh* gene that lacks the piRNA homologous sequences was cloned 5’ to the Luciferase gene. Thus, the use of antagopir to piRNA sequences present in UTR of the gene showed that piRNAs regulate gene expression.

#### 3.1.3 Deregulated genes identified by comparative sperm proteome analysis have UTR-homologies to the Y-transcripts

In a comparative sperm proteome analysis, using wild type (XY^RIII^) mice and a Y-deleted mutant mice (XY^RIII^qdel) we had previously identified a few proteins, calreticulin, FABP9, SOD and SPINK2V2 that were upregulated, SPINK2V3 that was downregulated and undetectable CFAP90, MAST and SDF2L. Therefore, these proteins, were differentially expressed between the wild type and the Y-deletion mutant [9]. UTRs of these genes had homologies with *Pirmy* and *Pirmy*-like RNAs. UTRs of these deregulated genes were also had homologous to the Y-transcripts being analyzed in this study. *Cyp19A1* upregulated in B10.BR-Ydel mice was also used for analysis [21]. Using BLAST the Y-transcripts found 16 short homologies. These homologies were found in the 3’UTRs of *Cfap90, Sdf2l1*, calreticulin, *Spink2V3* and *Cyp19A1* and the 5’UTR of *Fabp9*. Ten out of 16 homologous sequences were used as probes on northern blots andall of them identified signals corresponding to piRNAs (Figure 1). These probes sequences were searched against the mouse piRNA database, and a few of these regions did identify piRNAs. The probes used here are depicted on the UTRs of the deregulated genes (Figure S5). Thus, we identified six proteins that are deregulated in the sperms of the Y-deletion mutant mice, with piRNA motifs in their UTRs using the Y-transcripts.

### 3.2. MuRVY element in Orly1 has homology to UTRs of autosomal genes

The BLAST search of the 8 Y-transcripts against the mouse UTRs (UTRdb) based on the GRCm39 assembly using default parameters, yielded 27 hits larger than 150 nt. Two regions, one on *Orly1* [1303-1686 (384 nt)] and one on *Sly* [767-918 (152 nt)], were significant. A BLAST analysis of these two regions against the mouse genome revealed their presence in multiple chromosomes (Figure 3a, b). A reciprocal BLAST of UTRs (UTR db) against the transcripts yielded 36 hits, which included the 27 hits from the previous analysis. Of the 27 hits 5 were for *Orly1* and 22 for *Sly*.

**Figure 3.**
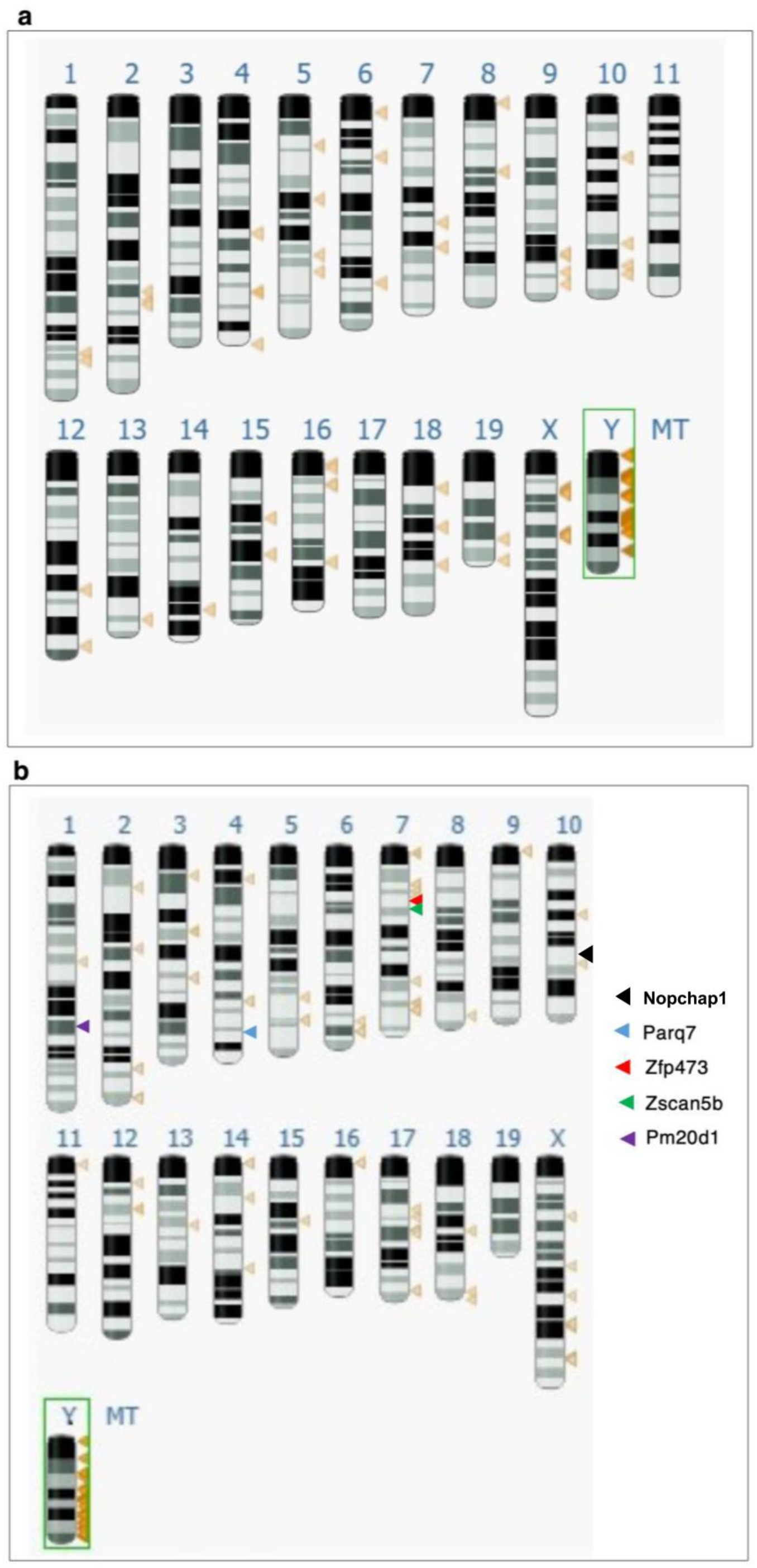
Mapping of the 152 nt-region of *Sly* **(a)** and 384 nt-region of *Orly1* **(b)** on the mouse genome are depicted as brown triangles. The intensity of the color indicates the quality of the match at each locus. The dark brown arrows correspond to stronger identities, while light brown arrows represent shorter hits or less identity. Genes which harbored the 384 nt sequence in their UTRs are marked in different colors as indicated within the figure.

All the genes, whose UTRs hit *Sly*, were located on the X chromosome (Figure 3a) and are *Sly* homologues, *Slx*. The *Sly-Slx* homologous stretch of 152 nucleotides identified 7 gold standard piRNAs. These 7 piRNAs hit a 40 nt region on *Sly* (544-583) and 39 nt-region on *Slx* (581-619). The regions on *Sly* and *Slx* are nearly identical, with just 3 mismatches. A dPCR analysis showed that there were approximately 734 copies of *Sly* in the wild type (XY^RIII^) genome whereas in the deletion mutant (XY^RIII^qdel) genome, the copy number average dropped to around 199 (Figure 4a). A several-fold increase in the expression was observed, for six of the *Sly* piRNAs analyzed in the study, in XY^RIII^qdel mice testes (Figure 4b).

**Figure 4.**
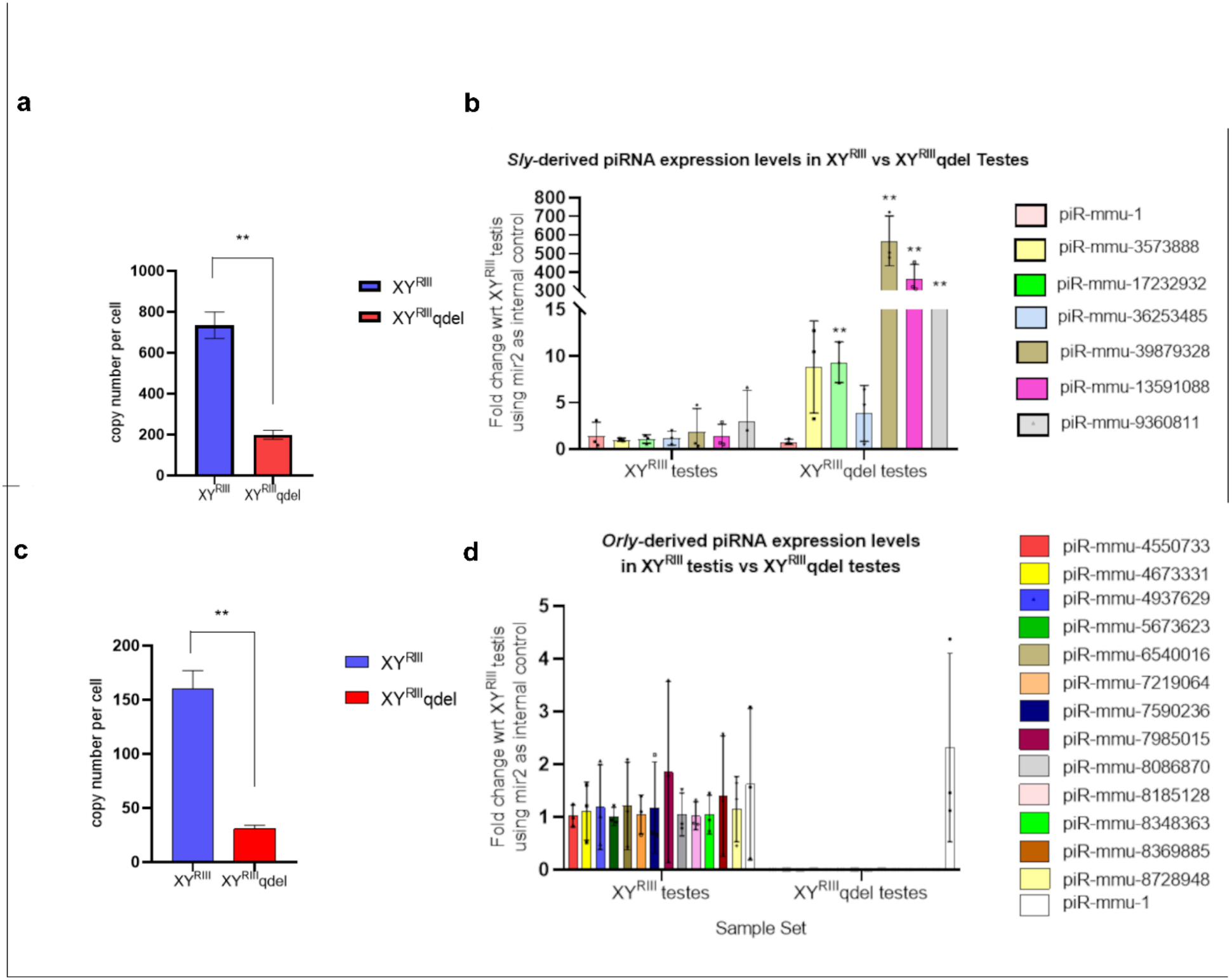
This Figure shows the copy number analysis (a, c) and expression profiling of XY^RIII^ and XY^RIII^qdel mice **(a)** histogram representing the copy number differences of *Sly* between the two genomes by dPCR analysis (Figure S6 e-h). **(b)** expression of piRNAs identified by *Sly* in testes of these mice (n=3; **= p≤0.01). **(c)** histogram representing the copy number difference of *Orly1* between these two mice by dPCR (Figure S6 a-d). **(d)** expression of piRNAs identified by *Orly1* in testes of these mice (n=3; significance was not represented as, expression in XY^RIII^qdel were not detectable by qPCR and Ct value was considered to be above 41 for graphical representation).

The 384 nt-region from *Orly1*, localized to its last exon which is not present in *Orly2* or *Orly3* and hence is specific to *Orly1*. The genes *Pm20d1, Paqr7, Zfp473, Zscan5b* and *Nopchap1*, whose UTRs matched *Orly1*, localize to the autosomes 1, 4, 7 and 10 (Figure 3b). When the 384 nt region was searched against the genomic sequences of these autosomal genes using the Ensembl Gene database, we found matches to UTR regions in all the 5 genes mentioned above. The 384 nt region was present in the 3’UTRs of *Nopchap1, Pm20d1, Paqr7* and 5’UTR of *Zfp473* and *Zscan5b* (Figure S6). Searching the repeat database DFAM (https://www.dfam.org) with *Orly1* revealed the 384 nt-region to be part of ERV1 LTR (Figure S6). To determine the number of copies of *Orly1* on Y chromosome, we searched for *Orly1* in the mouse genome by changing the default parameters (increasing the number of maximum matches in a query range to 100 and maximum target sequences to 5000). This analysis identified 102 copies. Shorter stretches of homologies were found on all the chromosomes (Figure 3b). A dPCR analysis showed that the number of *Orly1* copies in XY^RIII^qdel genome was on average 30 copies per cell compared to the average 160 copies per cell in XY^RIII^ mice (Figure 4c). A BLAST analysis of the 384 nt-region of *Orly1* identified 13 piRNAs in the piRNA database. The 13 piRNAs are clustered in two regions on *Orly1*, nine in 1368-1424 and four in 1568-1676. There was a drastic reduction in the expression of these 13 piRNAs in XY^RIII^qdel mice testes (Figure 4d).

We then studied the expression of PM20D1protein, coded by one of the four genes containing the ERV-homologous UTR. PM20D1 protein analyzed using antibody from Invtrogen (Cat no: PA5-46590), showed an approximately 1.8-fold higher expression in XY^RIII^qdel compared to its wild type XY^RIII^ (Figure 5a, b, Table S4).

**Figure 5.**
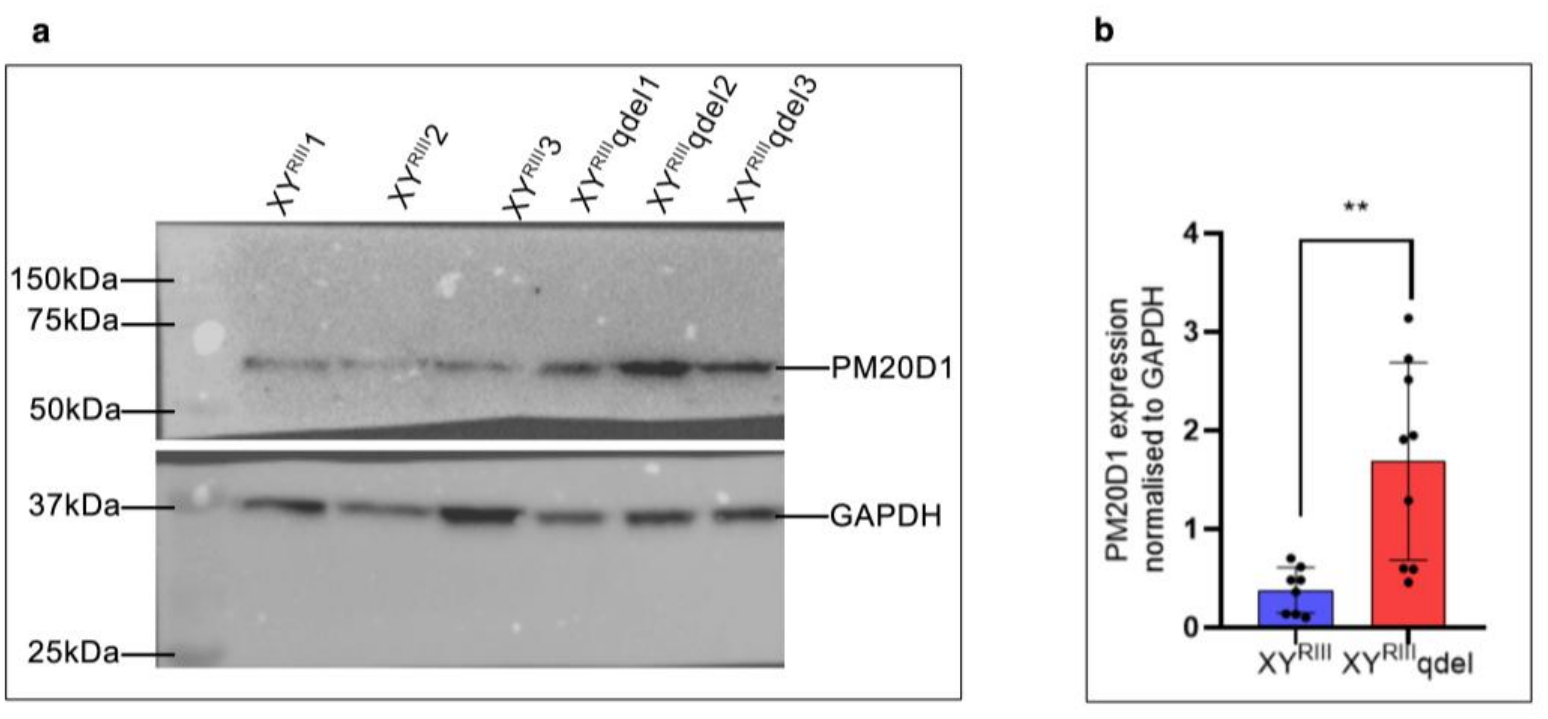
Comparison of PM20D1 expression in XY^RIII^ and XY^RIII^qdel testes. (**a**) Western blot image representing PM20D1 expression from 3 mice, each from XY^RIII^ and XY^RIII^qdel. Bottom panel represents signal from GAPDH (Thermo Fisher Cat no:MA5-27912) used as loading control. (**b**) Histogram representation of PM20D1 expression from data points (Table S4) including biological and technical replicates (n=9; **= p≤0.01).

Thus, along with the previous proteomics study in the lab, we have identified 10 proteins that have piRNA homologies in the UTRs of the corresponding genes. Seven of these proteins show differential expression in the proteomics screen and the western blot analysis put together.

### 3.3 Human homologues of the deregulated mouse genes identify human piRNAs

The human homologues of those genes identified as deregulated in XY^RIII^qdel mice in our previous study [9] were extracted from the NCBI database. UTRs of human homologues of Calreticulin, Spink V3 and *SOD* genes were BLASTed against the piRBase using default parameters; this identified human piRNAs; they do not identify mouse piRNAs.

## 4. Discussion

Based on the evidence presented here that (i) the identification of 30 nt RNAs mainly in testis, using Y-homologous sequences as probes present in UTRs of different genes (ii) differential expression from two DNA strands and (iii) identification of corresponding piRNAs in different piRNA databases we establish that mouse Y multicopy gene transcripts generate piRNAs. In addition, we have identified a 384 nt long MuRVY element, an LTR belonging to ERV (class I) homologues of which, are also present in the UTR regions of a few autosomal genes. These ERV homologous segments identify piRNAs. We also demonstrated upregulation of one of these genes in XY^RIII^qdel mice testes.

PIWI interacting RNAs (piRNAs) that range in size from 26-32 nt were reported initially from mouse testis by four different groups [19, 22-24]. piRNAs are generated from transposons and are involved not only in transposon silencing in but also genomes and germ cell maintenance [25-27]. piRNA generating clusters were reported from all the chromosomes except the Y in mammals [19, 24, 28, 29], except for a report of piRNAs from *Pirmy*, its splice variants and transcript variants on mouse Y chromosome [9]. The suppressor of stellate (*Ste*) locus in *Drosophila* is another example of a Y-chromosomal piRNA cluster [26, 27]. The presence of repeat sequences *Pirmy, Sly, Ssty1, Ssty2, Asty* and *Orly1* on the long arm of mouse Y chromosome generating piRNAs, confirms that the long arm Y-harbors piRNA generating sequences.

A couple of groups have reported piRNAs, having roles in spine development and long-term memory in brain of mouse and Aplysia [30, 31]. The observation of Y chromosome-derived piRNAs in mouse brain in the present study (Figure 1i), is commensurate with reports of gene expression from the Y chromosome in mouse brain [32]. An earlier study on the stress-induced behavior of XY^RIII^ and XY^RIII^qdel males showed altered neural gene expression in the latter [17]. These genes, which showed deregulated expression in brain, had homology to Y-derived noncoding RNAs in their UTRs, which suggests regulation of these genes by Y-derived piRNAs. Thus, we find that the mouse Y multicopy gene transcripts generate piRNAs for gene regulation in brain and testis.

Different groups have hypothesized a gene on the male-specific region of the mouse Y (MSYq) that could regulate X chromosomal gene expression [11, 12, 33]. Earlier work in our lab had demonstrated regulation of autosomal genes expressed in testis by piRNAs from multicopy Y-linked *Pirmy* [9]. In the present study we further observe regulation of autosomal genes by piRNAs derived from additional Y-chromosomal transcripts *Sly, Ssty1, Ssty2, Asty* and *Orly1*.

A partial, degenerate copy of MuRVY, belonging to the ERV class I is present in the *Orly1* transcript [8]. We propose that part of the LTR of this ERV found in the UTRs of the autosomal genes regulates theexpression of these genes, possibly through the piRNAs that are identified within this segment. All the thirteen piRNAs identified by *Orly1* are present within the LTR sequence. An accumulation of Transposable Elements and small RNAs have been reported in the 5’ and 3’ UTRs of genes, which have been associated with gene expression levels [34].

The increased number of *Sly*-homologous piRNAs on reduction of the copies of *Sly* on the Y-chromosome could perhaps be generated from the 40-nucleotide *Sly-Slx* homologous region on the X-chromosome. This would perhaps reflect a conflict between the X- and Y-chromosomes described earlier by several groups earlier [11,35]. *Slx* is reported to be overexpressed in XY^RIII^qdel testis, which shows a reduction in *Sly* copy number [11,36]. On the other hand, a deletion of the Y-exclusive *Orly* copies exhibits a reduction in expression of *Orly*-derived piRNAs. At the same time, an upregulation of PM20D1 expression in XY^RIII^qdel mice testis shows the effect of Y-chromosomal deletion on an autosomal gene that harbors the ERV-LTR from *Orly1* and the *Orly1*-derived piRNAs. Shipz et al. [37] have envisaged euchromatic target insertions as sites for production of pi- and endo-siRNAs in *Drosophila*. On the contrary, here we find a drastic reduction in the expression of *Orly1*-derived piRNAs with the deletion of the Y-chromosomal copies, even though these sequences are present in the UTRs of autosomal genes, implying that the piRNAs are produced by *Orly* repeats on the Y chromosome, not from autosomal genes. The homologous sequences present in the UTRs of the autosomal genes appear to be targets of these piRNAs. The elevated expression of PM20D1 protein in the Y-deleted (XY^RIII^qdel) mice is a clear example of such a regulation. Deregulated expression of autosomal genes with piRNA homologous sequences in the corresponding UTRs has been reported earlier in XY^RIII^qdel mice [9, 17].

Calreticulin, SPINK2, SOD, FABP9, CFAP90 and SDF2L1 are some of the proteins which were differentially expressed between the wild type and Y-deletion mutant mice in a comparative sperm proteomics study [9]. In the earlier study, we had opined that these genes are regulated by *Pirmy*-derived piRNAs. But the present study shows that the same genes are regulated by other Y-transcripts also. The identification of UTRs of deregulated genes by different Y-transcripts, elucidates the fact that each gene is regulated by multiple Y-transcripts. The UTRs of these, identified mouse piRNAs in piRBase database. The identification of piRNAs by the UTRs of human homologues of Calreticulin, *Spink2* and *Sod* broadens the relevance of the study with respect to humans. *SOD* is one of the markers for male infertility in humans [38]. *SPINK2* has also been shown to be associated with human male infertility [39]. As we identify piRNAs using UTRs of human homologues of these genes, Calreticulin, *Spink2* and *SOD*, it is plausible that similar mechanisms of piRNA mediated regulation could exist in humans as well.

The protein-coding genes, degenerate pseudogenes, and repeat sequences on the mouse Y-long arm are indispensable for normal spermatogenesis [18]. The up/down-regulation of gene expression from other chromosomes on deletion of the Y [12] suggests interaction between the Y chromosome and the X chromosome/autosomes. Current study provides further evidence in support of the physical connection between mouse Y chromosome and transcripts from the rest of the genome via piRNAs. The multicopy *Ssty, Sly, Asty* and *Orly1* genes located on the mouse Y chromosome, generate piRNAs that potentially regulate expression of X-chromosomal and autosomal genes. *Ssty* and *Sly* also code for proteins. It is possible that the noncoding *Sly, Ssty* transcripts [4, 10, 11,14] are used by the genome to generate piRNAs for gene regulation. Owing to the complexity and multicopy nature of these Y-chromosomal genes, it is difficult to differentiate between the coding and noncoding transcripts. Crosstalk between the Y-chromosome and the rest of the genome, both X- and autosomal, by genes on the Y-chromosome, appears to be a common theme of regulation in genomes.

This putative regulatory role of Y-chromosomal non-coding transcripts via piRNAs further strengthens the hypothesis of regulation of autosomal genes by noncoding transcripts from Y-heterochromatin repeats in testis [16, 9, 17], albeit through different mechanisms. A combinatorial approach of gene expression analyses using Y-deletion models together with characterization of noncoding RNAs from corresponding Y chromosomes may elucidate more examples of such interactions. The repeat sequences on Y-chromosomes appear to have been engineered for versatile roles.

## Acknowledgements

We wish to thank late L. Singh, former Director, Centre for Cellular and molecular Biology, Hyderabad for discussions, Jennifer Marshal Graves, Distinguished Professor at La Trobe University, Australia, S. C. Lakhotia, Distinguished Professor, Banaras Hindu University Varanasi, India and V. Radha, Scientist (retired), CCMB, Hyderabad for inputs during manuscript preparation and M. R. Sunayana and R. Nandini for help with the figures. Prof. Rajiva Raman, Professor, Banaras Hindu University, Varanasi, India is acknowledged for his kind help in critical reading and language editing of the manuscript.

## Author Contributions

RAJ - conceived the idea and directed the project, KM, AP - did the northern blots and other molecular biology experiments, AC, SK, ST - did in silico analysis, NMP - did the gel shifts, HMR-initiated the experiments, JLA did the functional analysis, AM and AK performed the piRNA PCR and dPCR experiments.

## Institutional Review Board Statement

Mice used in this study were bred in our animal house facility; we have ethical committee clearance from our Institutional Review Board to use the mice (IAEC 65/28/2006). Animal study protocols were approved by our institutional ethics committee.

