## Supplemental data for "piRNAs from Y chromosomal protein coding, noncoding and endogenous retrovirus homologous repeat families regulate autosomal gene expression in mouse testis"

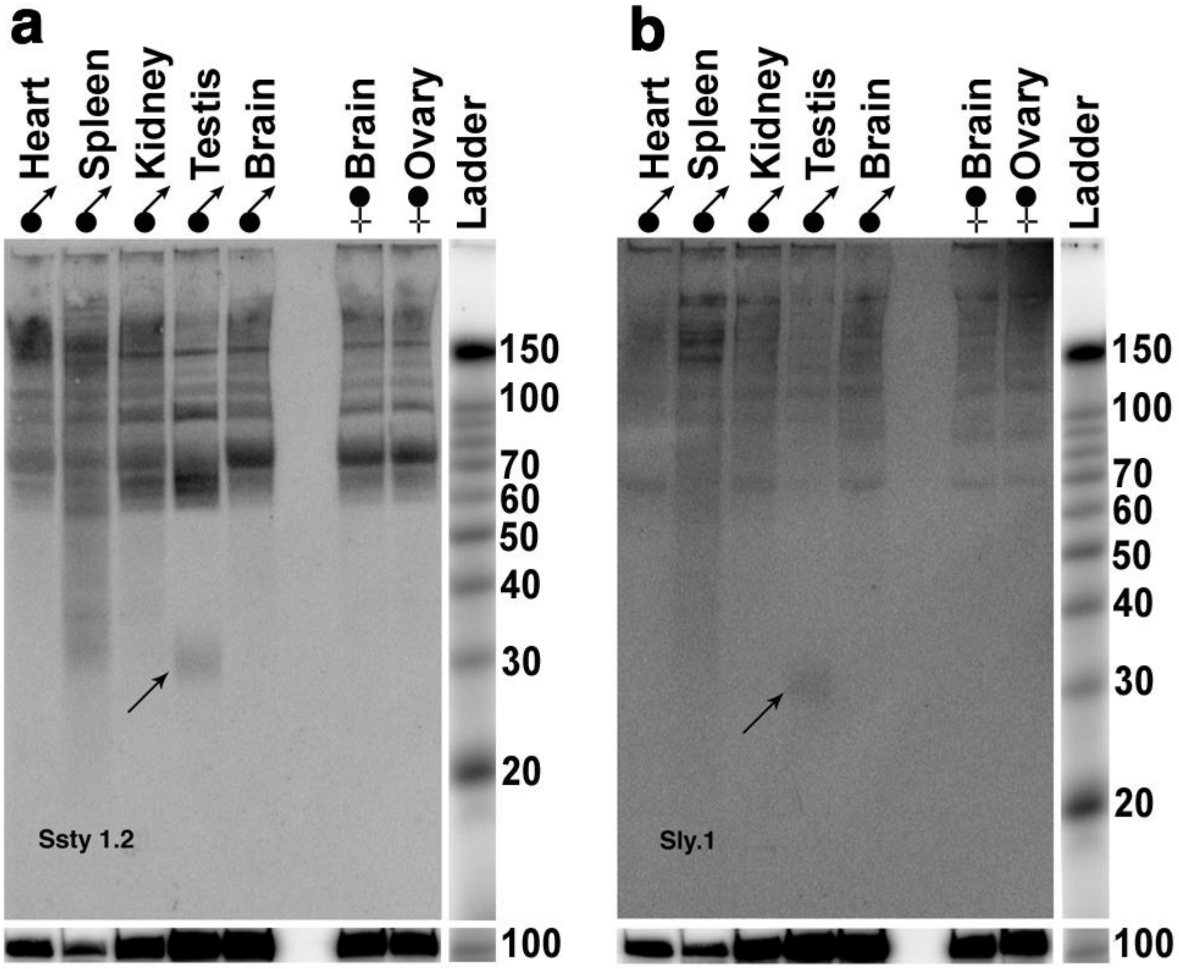


**Figure S1: Multi-tissue small RNA northern blot.** Northern blot including small RNA from male and female tissues (ovary and brain) showed ~30 nt signals in testis only, with (a) Ssty1.2 and (b) Sly.1 probes showing the specificity of the probes. U6 was the loading control for both the probes.

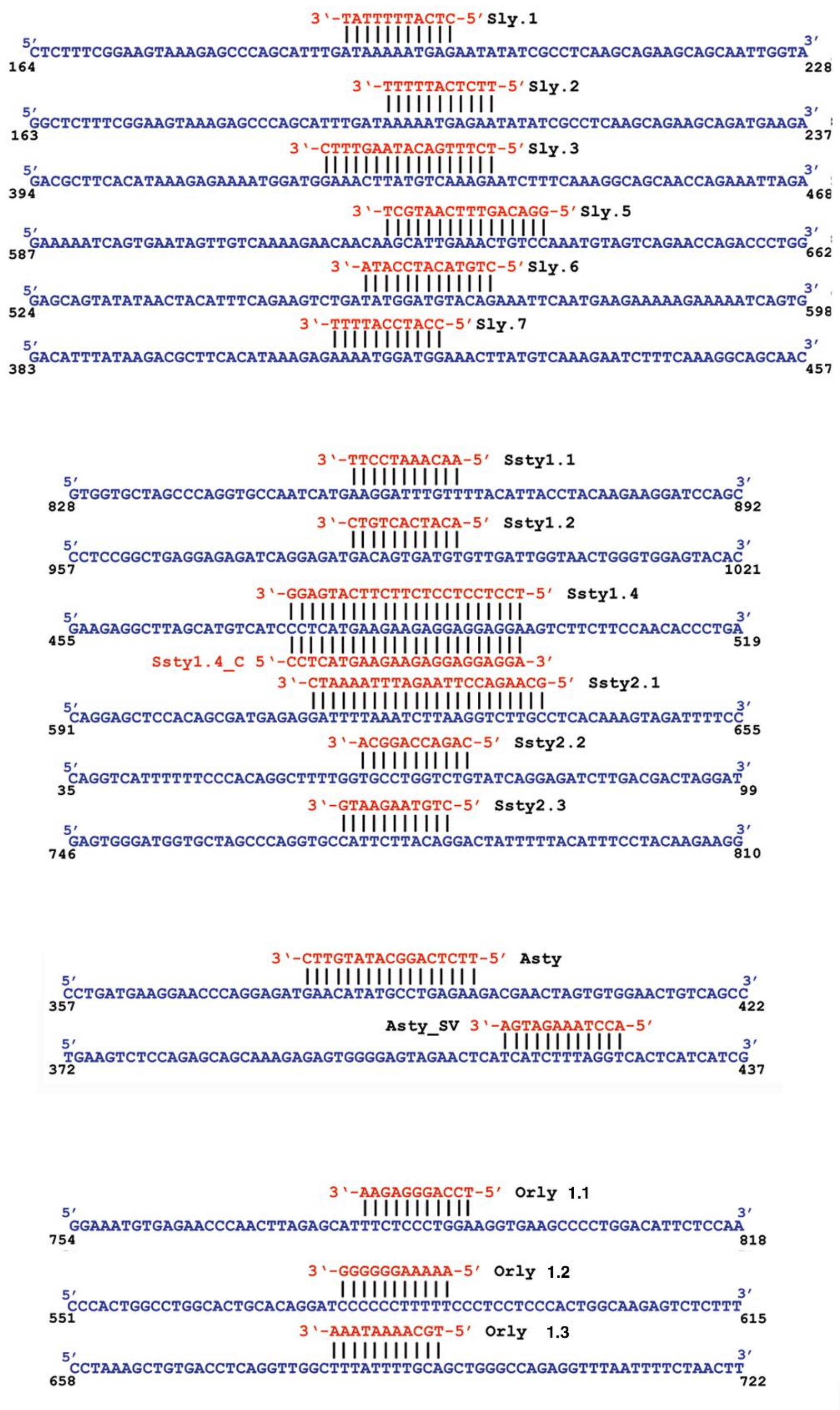

**Figure S2**: **Localization of probes to the Y-transcripts:** Positions of the small RNA probes (red colour) used for northern blotting on corresponding Y-transcripts (blue colour). Antisense probes are shown below the strand sequence with the corresponding gene names on the left.

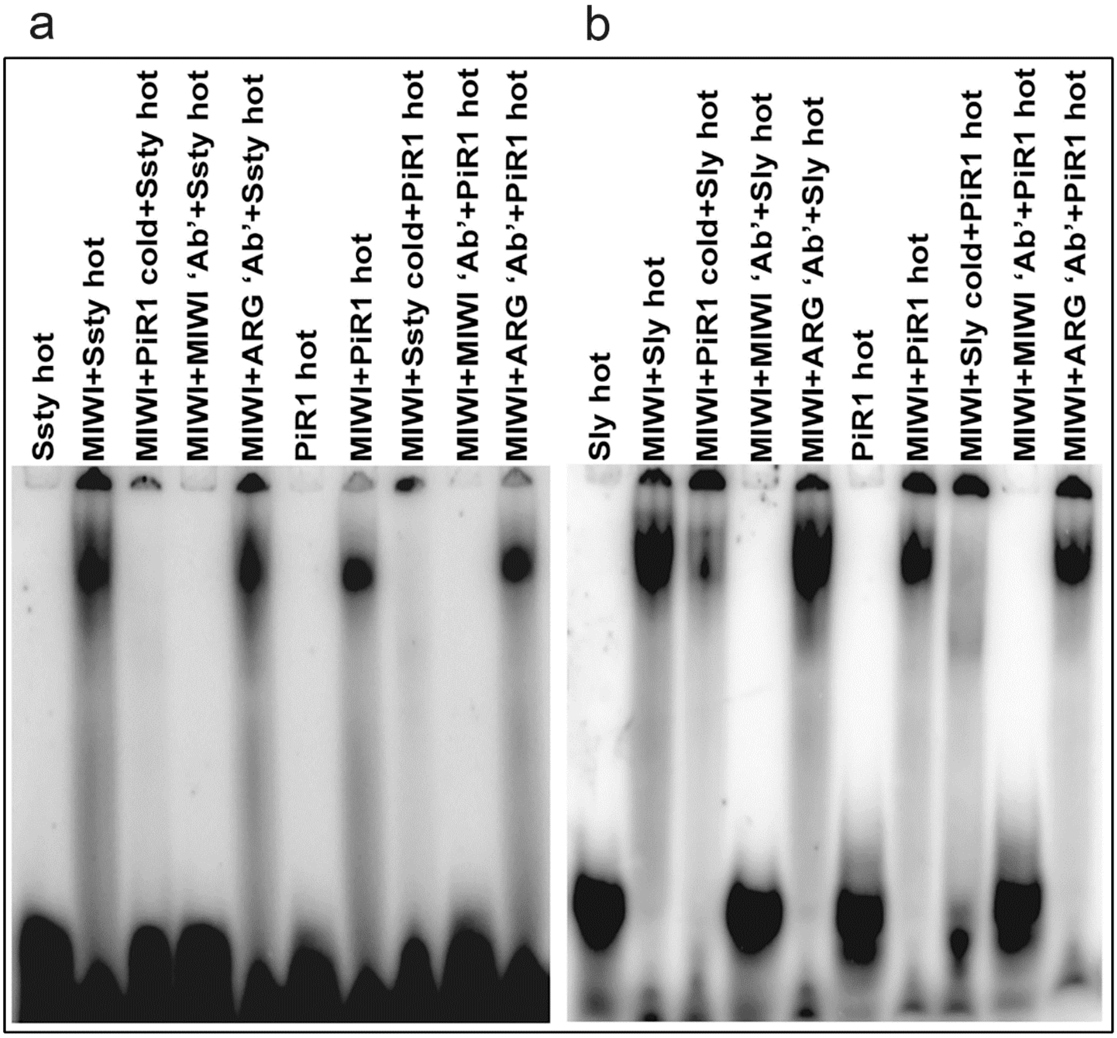


Figure S3. Putative piRNAs from mouse Y-chromosome bind to MIWI protein: EMSA with MIWI protein and putative piRNA oligonucleotides from (a) *Ssty*; (b) *Sly* showed shifted bands similar in size to that of piR-mmu-1; the gel-shifted bands were competed out on use of cold *Ssty*/*Sly*/piR-mmu-1 oligonucleotides and MIWI antibody. A nonspecific antibody, Argonaute 3, did not compete out binding of the putative piRNAs to *Ssty*/*Sly* or piR-mmu-1.

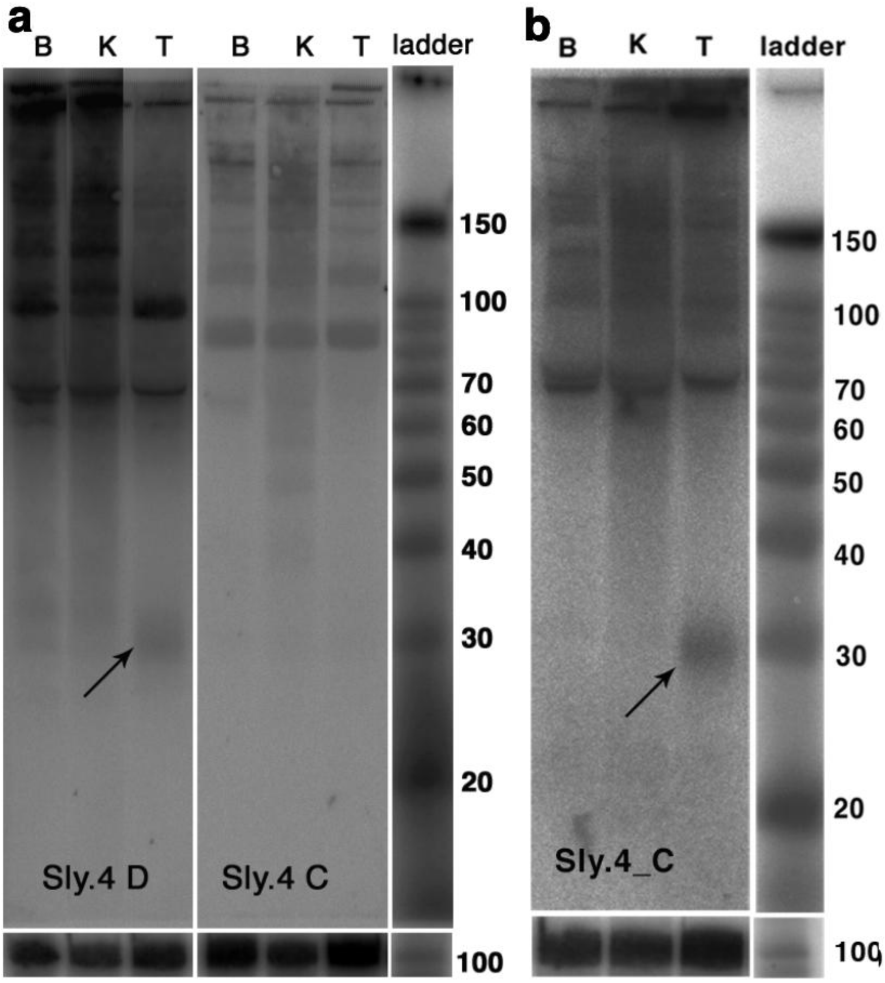


**Figure S4: Small RNA northern blots showing differential expression of piRNAs from direct and complementary strands.** Equal amounts of small RNAs from (B) brain, (K) kidney and (T) testis were electrophoretically separated, blotted, hybridized with probe noted on each blot, washed, developed and processed simultaneously. **(a, c)** LNA probes for predicted motifs from (D) direct and (C) complementary strands showed differential expression. **(b, d)** Blots hybridized with probes from complementary strands, revealed signals of approximately 30 nt in size, when washed under more stringent conditions and processed in the Sigmoid scale, indicating production of piRNAs from the complementary strand as well. U6 hybridization for each blot is given below the corresponding blot.

*Cfap90* 3'UTR

```
 CTGTAAGAATG (Ssty2.3)
 |||||||||||
5'ctgtaagaatgttctttgactgcaggaggataagcacgtgcatatctaaggtggtcctggctcagaacctggaa 3'
677                                                                         750
                    TTCCTAAACAA (Ssty1.1)
                    |||||||||||
5'ctcttcactttgcaaggatttgttggggaaggggtcacaggagaaacgtgccggtccttctccctgcccacaat 3'
897                                                                         970
               CTGTACATCCATA (Sly.6)  TCCAGGGAGAA (Orly1.1)
               |||||||||||||          |||||||||||
5'ttaaagccccctgtacatccatactgccaccccctccagggagaatgtgttacagattcagggcgcaggctggac
1082
                                                   AGTAGAAATCCA (Asty.sv)
                                                   ||||||||||||
  tctgggaaaagtctgcctgttacgttctctctctgtgatctgatgttgtgtcatctttaggtgaaatttctaat 3'
                                                                         1229
```

*Sdf2l1* 3'UTR

```
                                                           CTCATTTTTAT (Sly.1)
                                                           |||||||||||
5'tgagtgctggtattaaaggtgtacaccaccacaactggccccaacctgttcaataaactcattttttattactaa 3'
1036                                                                         1109
```

Calreticulin 3'UTR

```
5'ttcatttttccaggcaggttcagatctggggtagattctgatttgttcccctgcctcccccattacccc
1482
GGGGGGAAAAA (Orly1.2)                                    TTCTCATTTTT (Sly.2)
|||||||||||                                              |||||||||||
cccccctttttttttttactggtgtttgtctttaattctccttcagccctcatctggtttctcatttttgaatcaa 3
                                                                         1628
```

*FABP9* 5'UTR

```
                             ACGGACCAGAC (Ssty2.2)
                             |||||||||||
5'gcccacacttcatggttttcggttgtgaatgcctggtctgcccagaaagctgtgcacg 3'
1                                                          58
```

*Spink2* SV3 3'UTR

```
                         CTGTCACTACA (Ssty1.2)
                         |||||||||||
5'gaacttctctgggtagaggctggaagacagtgatgtaaagcggtttataaacttaaaactcttgttctgggctt 3'
486                                                                          559
```

*Cyp19A1* 3'UTR

```
                                                      CCATCCATTTT(Sly.7)
                                                      |||||||||||
5'agaagtgtggcctatctactctggggcattacttatccgtaattacatggaaaccatccattttatcgagtagg 3'
1623                                                                         1696
```

**Figure S5: Localization of the northern blot probes on to UTRs of deregulated genes:** The probes represented by matching lines in black are probes from the sense strand, and the red lines represent probes from the antisense strand.

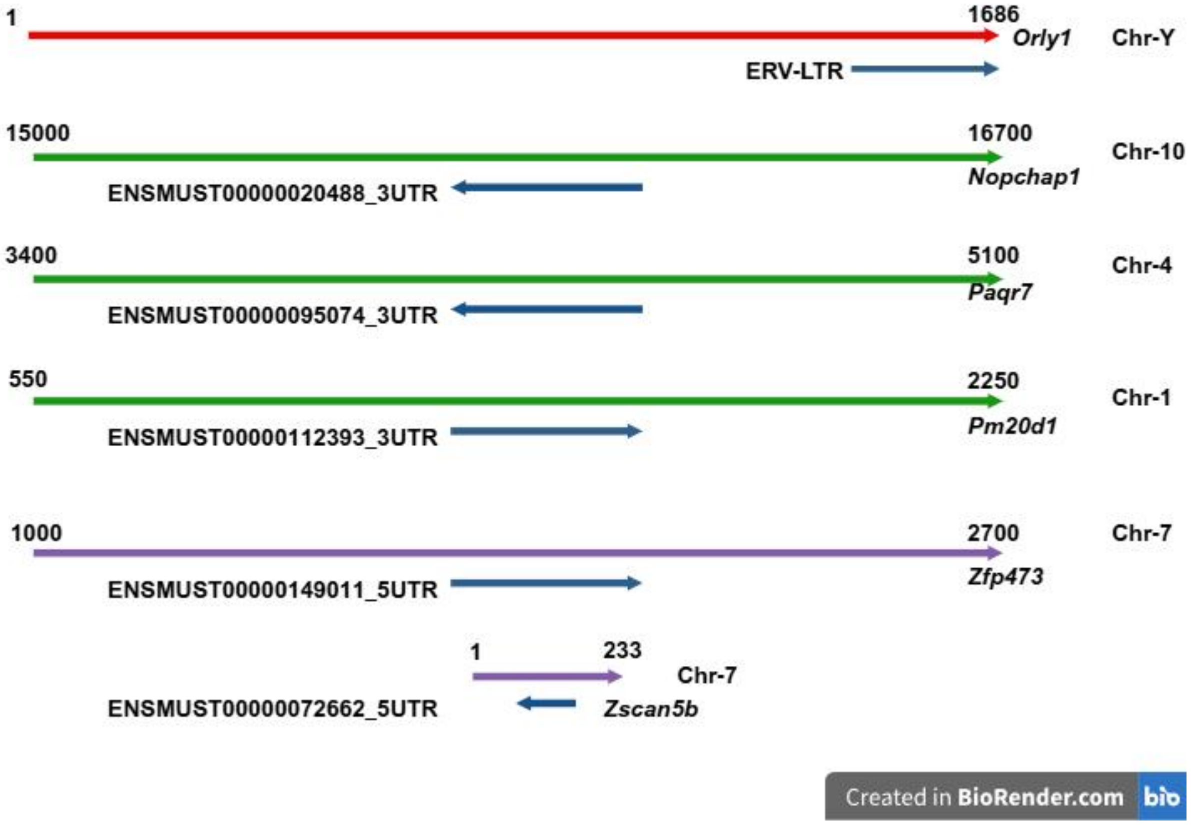


**Figure S6: Localization of the 384 nt homologous stretch on to autosomal UTRs.** Red arrow depicts the *Orly1* transcript and blue arrow the 384 nt stretch. Green and purple arrows indicate 3' and 5' UTRs of the autosomal genes respectively. Arrow heads on the ERV-homologous segment (blue arrows) represent the orientation of match within the UTRs. The entire Zscan5b 5' UTR (233 nt) and the short ERV-homologous match (80 nt) are shown. The other transcripts have longer UTRs, of which only 1700 nts have been depicted here.

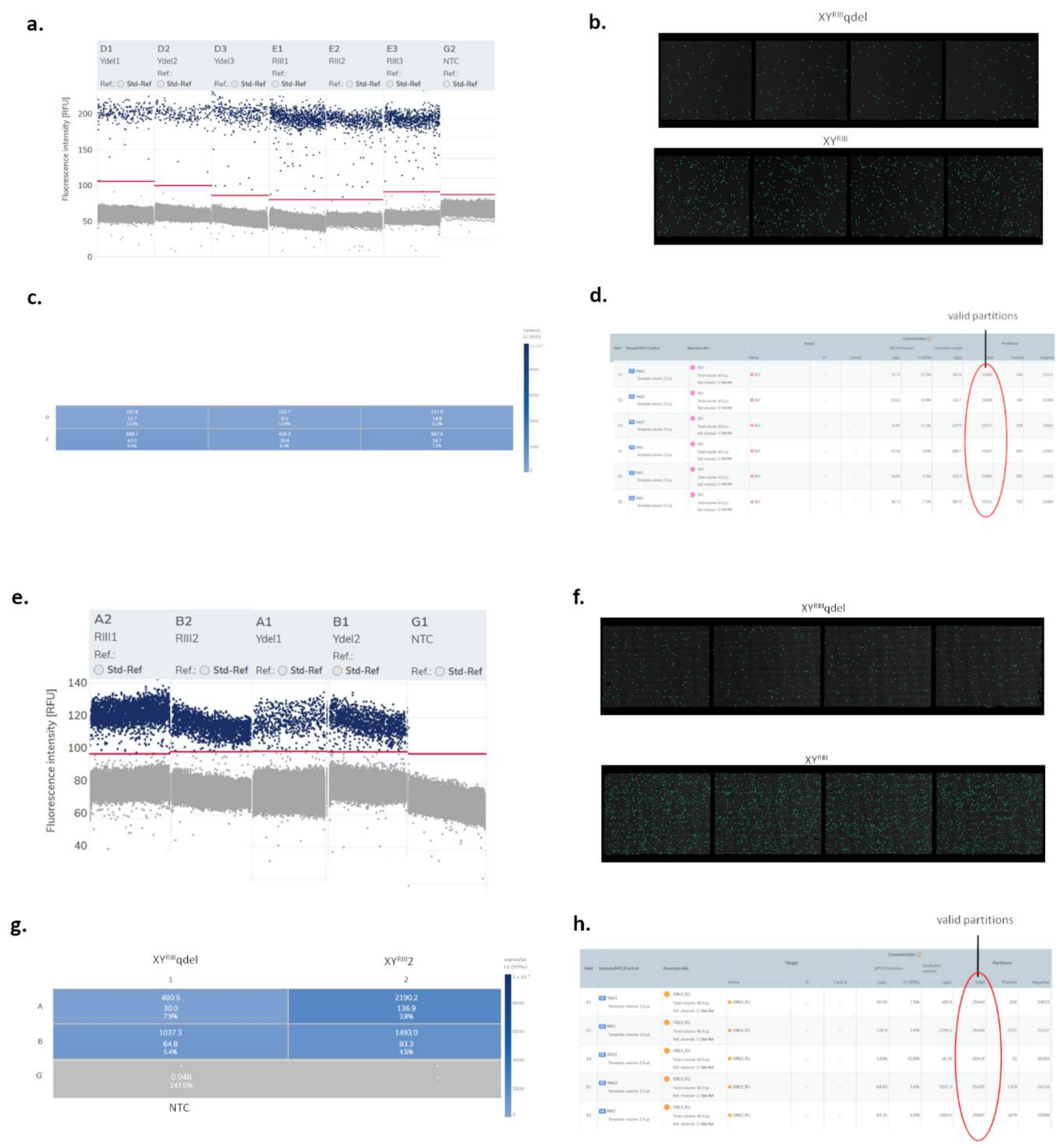


**Figure S7: Raw data images from dPCR experiment to estimate chromosomal copies of *Orly1* and *Sly*: (a)** one- dimensional 1d scatterplot showing dPCR signal density of *Orly1*in $XY^{RIII}$ (RIII1, RIII2, RIII3) vs $XY^{RIII}$qdel (Ydel1, Ydel2, Ydel3). **(b)** *Orly1* signal map showing the fluorescence detected by dPCR. **(c)** Heatmap showing relative differences of *Orly1* copies per ul between $XY^{RIII}$ and $XY^{RIII}$qdel. **(d)** Table showing copies detected per ul of reaction and the number of valid partitions considered for the experiment (valid partitions include partitions within the wells of the dPCR plate, that receives sufficient amount of sample for reaction validity, higher number of valid partitions ensure a high-fidelity experiment). **(e-h)** are the same depictions for *Sly*.

**Table S1**

| Y transcripts | Number of motifs in UTRs 3’ | 5’ | Number of motifs in either Orientation +/+ | +/- | Chromosome localization of target genes (*Mus musculus)* |
|---|---|---|---|---|---|
| *Sly* | 11 | 3 | 10 | 4 | 7 |
| *Ssty1* | 18 | 11 | 21 | 8 | 1, 7, 12 & 16 |
| *Ssty2* | 15 | 2 | 10 | 7 | 11 |
| *Asty* | 11 | 3 | 9 | 5 | 2, 3, 5, 9 & 17 |
| *Asty-sv* | 33 | 4 | 23 | 14 | 4, 9, 11, 12 & 13 |
| *Orly* | 11 | 1 | 5 | 7 | 8 |
| Total | 99 | 24 | 78 | 45 | - |

**Table S1. BLAST analysis of Y transcripts against UTRdb**: BLAST analysis of Y-transcripts against the mouse UTRs reveals short stretches of identity in the 5’ and 3’UTRs of genes localizing to different chromosomes in both +/+ and +/- orientations, indicating interactions between the Y-chromosomal and autosomal genes.

**Table S2**

| Probe | Sequence |
|---|---|
| Sly.1 | CTCATTTTTAT |
| Sly.2 | TTCTCATTTTT |
| Sly.3 | TCTTTGACATAAGTTTC |
| Sly.4 | TTCTCATGCATTTCTTTAACTG |
| Sly.4_C | CAGTTAAAGAAATGCATGAGAA |
| Sly.5 | GGACAGTTTCAATGCT |
| Sly.6 | CTGTACATCCATA |

| | |
|---|---|
| Sly.7 | CCATCCATTTT |
| Ssty1.1 | AACAAATCCTT |
| Ssty1.2 | ACATCACTGTC |
| Ssty1.4 | TCCTCCTCCTCTTCTTCATGAGG |
| Ssty2.1 | GCAAGACCTTAAGATTTAAAATC |
| Ssty2.2 | CAGACCAGGCA |
| Ssty2.3 | CTGTAAGAATG |
| Asty | TTCTCAGGCATATGTTC |
| Asty.sv | ACCTAAAGATGA |
| Orly1.1 | TCCAGGGAGAA |
| Orly1.2 | AAAAAGGGGGG |
| Orly1.3 | TGCAAAATAAA |
| Scr | GTGTAACACGTCTATACGCCCA |
| U6 | CACGAATTTGCGTGTCATCCTT |

**Table S2: Probes used in the study:** This table shows all the probes used in the study for northern blot analysis.

**Table S3**

| Query ID | Subject ID | Identity % | Alignment length | Query Length | Query Start | Query End | Subject Start | Subject End | E-Value | Bitscore |
|---|---|---|---|---|---|---|---|---|---|---|
| A1 | piR-mmu-60756543 | 100 | 31 | 622 | 374 | 404 | 31 | 1 | 1.43 E-07 | 61.9 |
| A1 | piR-mmu-53357055 | 100 | 31 | 622 | 296 | 326 | 1 | 31 | 1.43 E-07 | 61.9 |
| A1 | piR-mmu-60343372 | 100 | 30 | 622 | 232 | 261 | 1 | 30 | 5.66 E-07 | 60 |
| A1 | piR-mmu-31089132 | 100 | 30 | 622 | 266 | 295 | 30 | 1 | 5.66 E-07 | 60 |
| A1 | piR-mmu-17661661 | 100 | 30 | 622 | 142 | 171 | 1 | 30 | 5.66 E-07 | 60 |

| | | | | | | | | | | |
|---|---|---|---|---|---|---|---|---|---|---|
| A1 | piR-mmu-2923056 | 100 | 30 | 622 | 514 | 543 | 1 | 30 | 5.66 E-07 | 60 |
| A1 | piR-mmu-1882232 | 100 | 30 | 622 | 217 | 246 | 1 | 30 | 5.66 E-07 | 60 |
| A1 | piR-mmu-1692968 | 100 | 30 | 622 | 222 | 251 | 1 | 30 | 5.66 E-07 | 60 |
| A1 | piR-mmu-1612095 | 100 | 30 | 622 | 200 | 229 | 1 | 30 | 5.66 E-07 | 60 |
| A1 | piR-mmu-61183931 | 100 | 29 | 622 | 480 | 508 | 29 | 1 | 2.24 E-06 | 58 |
| A_Sv1 | piR-mmu-53357055 | 100 | 31 | 437 | 296 | 326 | 1 | 31 | 9.93 E-08 | 61.9 |
| A_Sv1 | piR-mmu-60343372 | 100 | 30 | 437 | 232 | 261 | 1 | 30 | 3.92 E-07 | 60 |
| A_Sv1 | piR-mmu-31089132 | 100 | 30 | 437 | 266 | 295 | 30 | 1 | 3.92 E-07 | 60 |
| A_Sv1 | piR-mmu-17661661 | 100 | 30 | 437 | 142 | 171 | 1 | 30 | 3.92 E-07 | 60 |
| A_Sv1 | piR-mmu-1882232 | 100 | 30 | 437 | 217 | 246 | 1 | 30 | 3.92 E-07 | 60 |
| A_Sv1 | piR-mmu-1692968 | 100 | 30 | 437 | 222 | 251 | 1 | 30 | 3.92 E-07 | 60 |
| A_Sv1 | piR-mmu-1612095 | 100 | 30 | 437 | 200 | 229 | 1 | 30 | 3.92 E-07 | 60 |
| A_Sv1 | piR-mmu-51919465 | 96.97 | 33 | 437 | 195 | 227 | 1 | 33 | 1.55 E-06 | 58 |
| A_Sv1 | piR-mmu-51776342 | 96.875 | 32 | 437 | 374 | 405 | 1 | 32 | 6.13 E-06 | 56 |
| A_Sv1 | piR-mmu-1955720 | 100 | 28 | 437 | 285 | 312 | 1 | 28 | 6.13 E-06 | 56 |
| Oy1 | piR-mmu-67173518 | 100 | 32 | 1137 | 907 | 938 | 1 | 32 | 6.73 E-08 | 63.9 |
| Oy1 | piR-mmu-53776028 | 100 | 32 | 1137 | 930 | 961 | 32 | 1 | 6.73 E-08 | 63.9 |
| Oy1 | piR-mmu-67293647 | 100 | 31 | 1137 | 778 | 808 | 31 | 1 | 2.66 E-07 | 61.9 |
| Oy1 | piR-mmu-67151422 | 100 | 31 | 1137 | 1072 | 1102 | 31 | 1 | 2.66 E-07 | 61.9 |
| Oy1 | piR-mmu-58378440 | 100 | 31 | 1137 | 595 | 625 | 31 | 1 | 2.66 E-07 | 61.9 |
| Oy1 | piR-mmu-51667752 | 100 | 31 | 1137 | 894 | 924 | 31 | 1 | 2.66 E-07 | 61.9 |
| Oy1 | piR-mmu-45606272 | 100 | 31 | 1137 | 831 | 861 | 1 | 31 | 2.66 E-07 | 61.9 |
| Oy1 | piR-mmu-67046246 | 100 | 30 | 1137 | 788 | 817 | 1 | 30 | 1.05 E-06 | 60 |
| Oy1 | piR-mmu-45587668 | 100 | 30 | 1137 | 304 | 333 | 1 | 30 | 1.05 E-06 | 60 |
| Oy1 | piR-mmu-45256536 | 100 | 30 | 1137 | 548 | 577 | 1 | 30 | 1.05 E-06 | 60 |

| | | | | | | | | | | |
|---|---|---|---|---|---|---|---|---|---|---|
| S1 | piR-mmu-51896273 | 100 | 33 | 1285 | 245 | 277 | 1 | 33 | 1.93 E-08 | 65.9 |
| S1 | piR-mmu-50685979 | 100 | 33 | 1285 | 350 | 382 | 1 | 33 | 1.93 E-08 | 65.9 |
| S1 | piR-mmu-3087745 | 100 | 33 | 1285 | 830 | 862 | 1 | 33 | 1.93 E-08 | 65.9 |
| S1 | piR-mmu-66013175 | 100 | 31 | 1285 | 861 | 891 | 1 | 31 | 3.01 E-07 | 61.9 |
| S1 | piR-mmu-64151258 | 100 | 31 | 1285 | 1007 | 1037 | 1 | 31 | 3.01 E-07 | 61.9 |
| S1 | piR-mmu-63268930 | 100 | 31 | 1285 | 620 | 650 | 1 | 31 | 3.01 E-07 | 61.9 |
| S1 | piR-mmu-53604364 | 100 | 31 | 1285 | 1216 | 1246 | 1 | 31 | 3.01 E-07 | 61.9 |
| S1 | piR-mmu-51971714 | 100 | 31 | 1285 | 1064 | 1094 | 1 | 31 | 3.01 E-07 | 61.9 |
| S1 | piR-mmu-1675809 | 100 | 31 | 1285 | 534 | 564 | 1 | 31 | 3.01 E-07 | 61.9 |
| S1 | piR-mmu-1441618 | 100 | 31 | 1285 | 609 | 639 | 1 | 31 | 3.01 E-07 | 61.9 |
| S2 | piR-mmu-1399949 | 100 | 34 | 1215 | 781 | 814 | 1 | 34 | 4.61 E-09 | 67.9 |
| S2 | piR-mmu-54114432 | 100 | 33 | 1215 | 851 | 883 | 1 | 33 | 1.82 E-08 | 65.9 |
| S2 | piR-mmu-52783168 | 100 | 33 | 1215 | 742 | 774 | 1 | 33 | 1.82 E-08 | 65.9 |
| S2 | piR-mmu-52534284 | 100 | 33 | 1215 | 1093 | 1125 | 1 | 33 | 1.82 E-08 | 65.9 |
| S2 | piR-mmu-52523219 | 100 | 33 | 1215 | 859 | 891 | 1 | 33 | 1.82 E-08 | 65.9 |
| S2 | piR-mmu-2631774 | 100 | 33 | 1215 | 1145 | 1177 | 1 | 33 | 1.82 E-08 | 65.9 |
| S2 | piR-mmu-335452 | 100 | 33 | 1215 | 781 | 813 | 1 | 33 | 1.82 E-08 | 65.9 |
| S2 | piR-mmu-54124931 | 100 | 32 | 1215 | 135 | 166 | 1 | 32 | 7.20 E-08 | 63.9 |
| S2 | piR-mmu-53786465 | 100 | 32 | 1215 | 1097 | 1128 | 1 | 32 | 7.20 E-08 | 63.9 |
| S2 | piR-mmu-53320913 | 100 | 32 | 1215 | 754 | 785 | 1 | 32 | 7.20 E-08 | 63.9 |
| Sy1 | piR-mmu-3461722 | 100 | 32 | 915 | 533 | 564 | 1 | 32 | 5.39 E-08 | 63.9 |
| Sy1 | piR-mmu-2402892 | 100 | 32 | 915 | 115 | 146 | 1 | 32 | 5.39 E-08 | 63.9 |
| Sy1 | piR-mmu-2093536 | 100 | 32 | 915 | 408 | 439 | 1 | 32 | 5.39 E-08 | 63.9 |
| Sy1 | piR-mmu-1483209 | 100 | 32 | 915 | 245 | 276 | 1 | 32 | 5.39 E-08 | 63.9 |
| Sy1 | piR-mmu-1483209 | 96.667 | 30 | 915 | 348 | 377 | 2 | 31 | 2.05 E-04 | 52 |

| | | | | | | | | | | |
|---|---|---|---|---|---|---|---|---|---|---|
| **Sy1** | **piR-mmu-4355014** | **100** | **31** | **915** | **379** | **409** | **1** | **31** | **2.13 E-07** | **61.9** |
| **Sy1** | **piR-mmu-4290171** | **100** | **31** | **915** | **146** | **176** | **1** | **31** | **2.13 E-07** | **61.9** |
| **Sy1** | **piR-mmu-3496674** | **100** | **31** | **915** | **178** | **208** | **1** | **31** | **2.13 E-07** | **61.9** |
| **Sy1** | **piR-mmu-2791755** | **100** | **31** | **915** | **77** | **107** | **1** | **31** | **2.13 E-07** | **61.9** |
| **Sy1** | **piR-mmu-2540676** | **100** | **31** | **915** | **349** | **379** | **1** | **31** | **2.13 E-07** | **61.9** |
| **Sy1** | **piR-mmu-2540676** | **96.552** | **29** | **915** | **247** | **275** | **1** | **29** | **8.11 E-04** | **50.1** |
| **Sy1** | **piR-mmu-2402891** | **100** | **31** | **915** | **115** | **145** | **1** | **31** | **2.13 E-07** | **61.9** |

**Table S3. Blast results of the Y transcripts (Sly, Ssty1, Ssty2, Asty, Asty-sv, Orly1) against the gold standard piRNAs downloaded from piRBase (2021). The abbreviations used are Sy1-Sly1; S1-Ssty1; S2-Ssty2; A1-Asty1; A_Sv1-Asty-sv, Oy1-Orly1;**

**Table S4**

| $XY^{RIII}$ | $XY^{RIII}qdel$ |
|---|---|
| 0.134537 | 0.599189 |
| 0.481913 | 0.592444 |
| 0.361941 | 0.458283 |
| 0.700202 | 1.90841 |
| 0.614269 | 3.139502 |
| 0.139836 | 2.51622 |
| 0.100124 | 1.949141 |
| 0.482437 | 2.726543 |
| | 1.289938 |

**Table S4**. **PM20D1 expression values used for Figure 6:** PM20D1 expression values were calculated from mean grey value relative to GAPDH for histogram in Figure 6.
